# Mechanistic Insights into Periplasmic Chaperones as Regulators of Protein Folding and Translocation in Gram-Negative Bacteria

**DOI:** 10.1101/2024.12.13.628315

**Authors:** Deep Chaudhuri, Madhu Bhatt, Shubhasis Haldar

## Abstract

In Gram-negative bacteria, periplasmic chaperones mediate the transport and folding of outer membrane proteins. While their substrate interaction mechanisms are well studied, their functional cycle during periplasmic translocation remains unclear. Using single-molecule force spectroscopy, we demonstrate that SecYEG-associated chaperones PpiD and DsbC act as foldases, delivering up to 45.5 zJ of mechanical work. This activity facilitates substrate extraction from the SecYEG translocation tunnel, reducing ATP consumption during SecA-mediated translocation. In contrast, chaperones such as Spy and Skp function as holdases, stabilizing unfolded substrates to prevent misfolding during translocation. Our findings reveal a unique mechanism by which chaperones adapt to mechanical constraints, optimizing protein folding and translocation in the periplasm.

## Introduction

Force plays a pivotal role in various biological systems, exerting its influence on numerous processes including cellular physiology, muscle construction, cell traction, protein folding, and translocation across the membrane^1–6^. Notably, in the context of protein folding, these force stimuli could regulate important biochemical signals through mechanotransduction and drive several post-translational translocation processes in cellular regions^7–10^. For instance, in bacterial cells, SecA has been demonstrated to generate a mechanical force of 10 pN at the Sec YEG tunnel allowing the translocation of the substrate proteins through the tunnel^10^. Additionally, co-translation folding at the ribosome’s entrance also takes place under mechanical force conditions^11^. These forces are instrumental in initiating the translocation of the secretory proteins^12,13^. Molecular chaperones, crucial actors in the heterogeneous protein folding process, are well known to interact with the substrate protein under mechanical stress, particularly in physiological contexts^14–17^. The interaction between the substrate and the chaperones constitutes a highly intricate and dynamic phenomenon, which is challenging for conventional experimental methodologies based on bulk techniques. Recent advancements in single-molecule techniques have a significant ability to examine individual molecular entities which has contributed to unrevealing the molecular mechanism of the chaperones^4,18^. These approaches have illuminated the complex roles of ATP independent periplasmic chaperones in protein folding and translocation across the membrane, uncovering their intricate molecular mechanism^19,20^. For example, an atomic force microscopy (AFM) study by Thoma et al. shed light on how SurA and Skp are influencing the folding pathway and modulating the free energy landscape during the membrane protein folding of FhoA^19^. More recent investigations, using fluorescence correlation spectroscopy (FCS) and smFRET, have enhanced our understanding of the roles of periplasmic chaperone SurA and Skp both as holdase and disaggregates for outer membrane proteins (OMPs)^21,22^. Despite these existing studies on the periplasmic chaperones in the folding process of the outer membrane protein, there remains a critical knowledge gap concerning their responses to mechanical stress.

The implementation of the single molecule force spectroscopy technique provides us with a powerful approach to characterize the change in the folding dynamics and molecular mechanism of a single protein molecule having precise spatial and temporal resolution^14,23–25^. In the context of a single molecule approach, the force can be utilized as a denaturant where the chaperone remains unperturbed, and force is exclusively applied to the client protein. In this study, we have monitored the mechanical role of periplasmic chaperones with our custom-made single-molecule covalent magnetic tweezers having the advantage of force ramp and force clamp techniques^14,23,24^. The force clamp experiment enables us to investigate the kinetic and thermodynamic properties of the client protein. The advantage of a long force range of 1-120 pN allows magnetic tweezers to detect the chaperone protein interaction at both folded, unfolded, and equilibrium states at sub-pN resolution. Moreover, the mechanical strength of the protein can be obtained by the force ramp technology by applying the force at a constant velocity with minimal thermal fluctuation. Collectively, these advancements allow us to detect how periplasmic chaperones could perturb their role in the presence of mechanical load^14^.

In this study, we conduct a direct measurement of mechanical force-induced conformational change and the folding dynamics of chaperone-protein interaction, which has uncovered the force-dependent role of the periplasmic chaperone in protein translocation across the periplasm. Here, our study focused on the globular B1 domain of protein L and the R3 IVVI domain of talin, both widely recognized as a model substrate^16,26–28^. Protein L is a two-state model protein synthesized in the cytosol and traversed over the periplasm, necessitates interaction with the periplasmic chaperones^25^. Here, we have explored the effect of PpiD and DsbC, associated with the SecYEG tunnel, along with other two periplasmic chaperones Skp and Spy, that could potentially interact with the substrate during translocation across the periplasm^29–32^. Initially, we focused on the tunnel-associated periplasmic chaperone, showing foldase activity under mechanical load, expanding additional mechanical work to facilitate protein folding. This additional mechanical energy generates a pulling force that reduces the ATP consumption by SecA during the substrate translocation towards the periplasm. Subsequently, we extended our findings to investigate another two periplasmic chaperone Skp, involved in OMP biogenesis, and Spy, a flexible periplasmic chaperone both acts as foldase as well as holdase^19,33^. Our observations indicate that both Spy and Skp act as holdase, operating effectively both in the presence and absence of mechanical force. Therefore, they hindered the folding process to prevent misfolding of the substrate and facilitating the translocation of substrate proteins from the periplasm to the outer membrane. To extend the applicability of our observations, we expanded our experiments with structurally different mechanosensitive protein talin, which has biological relevance in the chaperone interactions under force conditions^23,26,27^. Our result highlights a distinct force-driven mechanism employed by tunnel-associated periplasmic chaperones (PpiD and DsbC), which promote refolding by the utilization of binding energy to extract proteins from the SecYEG tunnel, while others (Skp and Spy) chaperones act as holdase, stabilizing the unfolded state. Altogether, these data describe a core mechanism of how the interplay of mechanical force can regulate chaperone action and facilitate the binding, folding, and release of the substrate towards the outer membrane.

## Results

### Folding dynamics of Single protein molecule measured by covalent magnetic tweezers

To study the folding dynamics of protein, we engineered a polyprotein construct of protein L comprising eight identical domains with Avi tag at the C terminus and Halo tag at the N terminus. One end of the protein L octamer is anchored to a glass surface via Halo tag covalent chemistry and another end is attached to a streptavidin coated paramagnetic bead (Fig. 1A)^14,24,25,34,35^. Force is applied on the paramagnetic bead to stretch the polyprotein construct by a pair of permanent magnets. The applied force can be adjustable up to sub pico-Newton range by changing the distance of the magnet to paramagnetic beads. The details of force calibration method have been discussed in the previous studies^25,14,24^. Our single molecule force spectroscopy technique allows us to study force-clamp experiments where we can unfold and refold the polyprotein construct by applying a constant force. Fig. 1B shows the representative stretching of the polyprotein at 45 pN force to observe eight district unfolding steps having the step size of about15 nm originating from protein L as a fingerprint. After complete unfolding, the force was decreased down to 7 pN and the polyprotein collapsed and experienced an instantaneous elastic recoiling followed by a discrete stepwise contraction. The number of unfolding steps precisely corresponds to the number of folded domains upon the application of a probe pulse. Since protein L do not form misfolded states, no significant differences in step sizes were observed in the presence of different chaperones (Supp. Fig. 1)^25,36^. Following the refolding process, the polyprotein reaches in the equilibrium phase where domains show folding-unfolding dynamics in between folded and unfolded state. The folding kinetics can be obtained as the first passage time (FPT) of unfolding and refolding i.e., the total time takes to unfold and refold the polyprotein completely. By averaging several trajectories mean first passage time (MFPT) of unfolding and refolding has been calculated from FPT^14,16,37,38^. The folding probability can be characterized from the equilibrium phase by dwell time analysis. The folding percentage of protein strongly depends on the applied refolding force, for instance folding probability of protein L is close to 1 at 4 pN and it shifts to 0 at 12 pN^16,34,39^.

**Figure 1:**
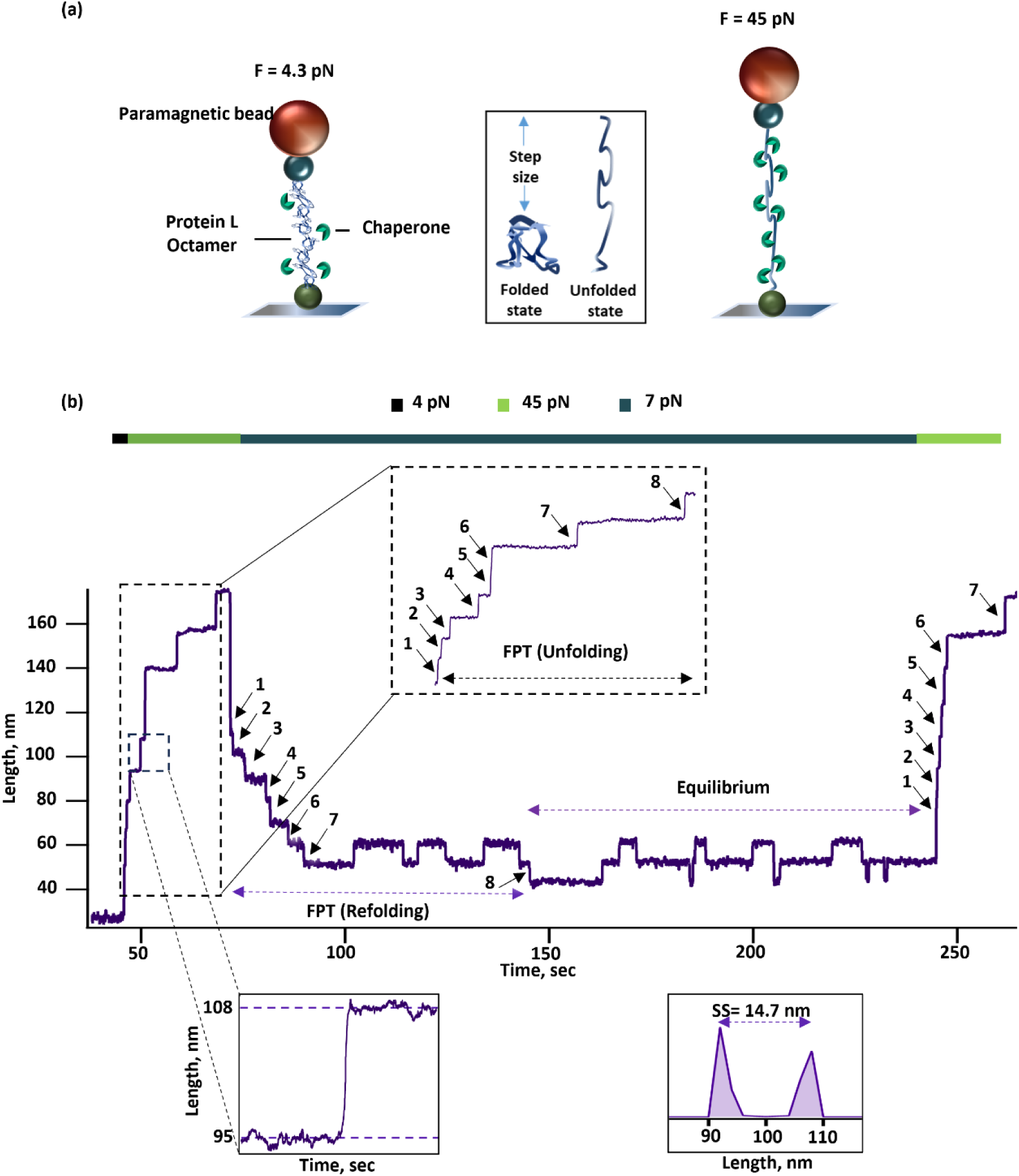
Experimental setup for magnetic tweezers experiments to study the folding dynamics of chaperone-protein interaction: **(A)** In the schematic depiction of single-molecule magnetic tweezers set up, the protein L octamer is tethered in between the streptavidin-coated paramagnetic bead and glass surface through Halo tag covalent chemistry. The applied force can be precisely controlled by changing the distance between the glass surface and the permanent magnet. **(B)** A representative trace of force-clamp experiment demonstrates that protein L can undergo complete unfolding at 45 pN, showing eight unfolding steps as fingerprints with a step size of 1̴ 5 nm. Quantitatively, a quenching pulse of 7 pN refolds the polyprotein completely after an elastic recoiling. Subsequently, the protein undergoes an equilibrium phase where it starts hopping in between the folded and unfolded state. The FPT (first passage time) of unfolding can be calculated as the total time to take the protein to unfold completely and the FPT of refolding is the total time to refold completely can be obtained from a single trace. Raw data was recorded and smoothed by using box algorithm. Step size was calculated by taking the distance between the two centres of the histogram plots.

### Mechanical foldase activity of tunnel-associated periplasmic chaperones

It is well known that chaperones can interact with the substrate at unfolded, partially folded, near native, and molten globule states to help in protein translocation which could be modulated by mechanical force^17,40^. PpiD, an ATP-independent multidomain periplasmic chaperone, associated with the SecYEG complex, interacts early with the newly synthesized substrate protein for translocation^32,41^. It has an important role in directly influencing the translocation efficiency of the SecYEG translocon by interacting with the substrate protein^29,32,41^. To further explore its role in protein folding, we sought to investigate the effect of PpiD on the folding dynamics of substrate protein under force. We focused on a concentration where the mechanical effect of PpiD reached saturation (Supp. Fig. 2), enabling us to evaluate its chaperone activity under force. We have measured the folding probability of protein L at varying forces within the range of 4- 12 pN under the influence of 25 µM PpiD (Fig. 2A). This comparison revealed that the FP of protein L dramatically increased in the presence of PpiD in a particular force range of 6 to 8 pN. The increment in the half-point force (force at FP 0.5) has been observed from 7.01 to ∼8 pN in the presence of PpiD suggesting the mechanical effect of PpiD under force. The maximum change in the folding probability has been observed at a particular force of 8 pN where the FP increased drastically in the presence of PpiD from ̴ 0.18 to ̴ 0.7 (Fig. 2B).

**Figure 2:**
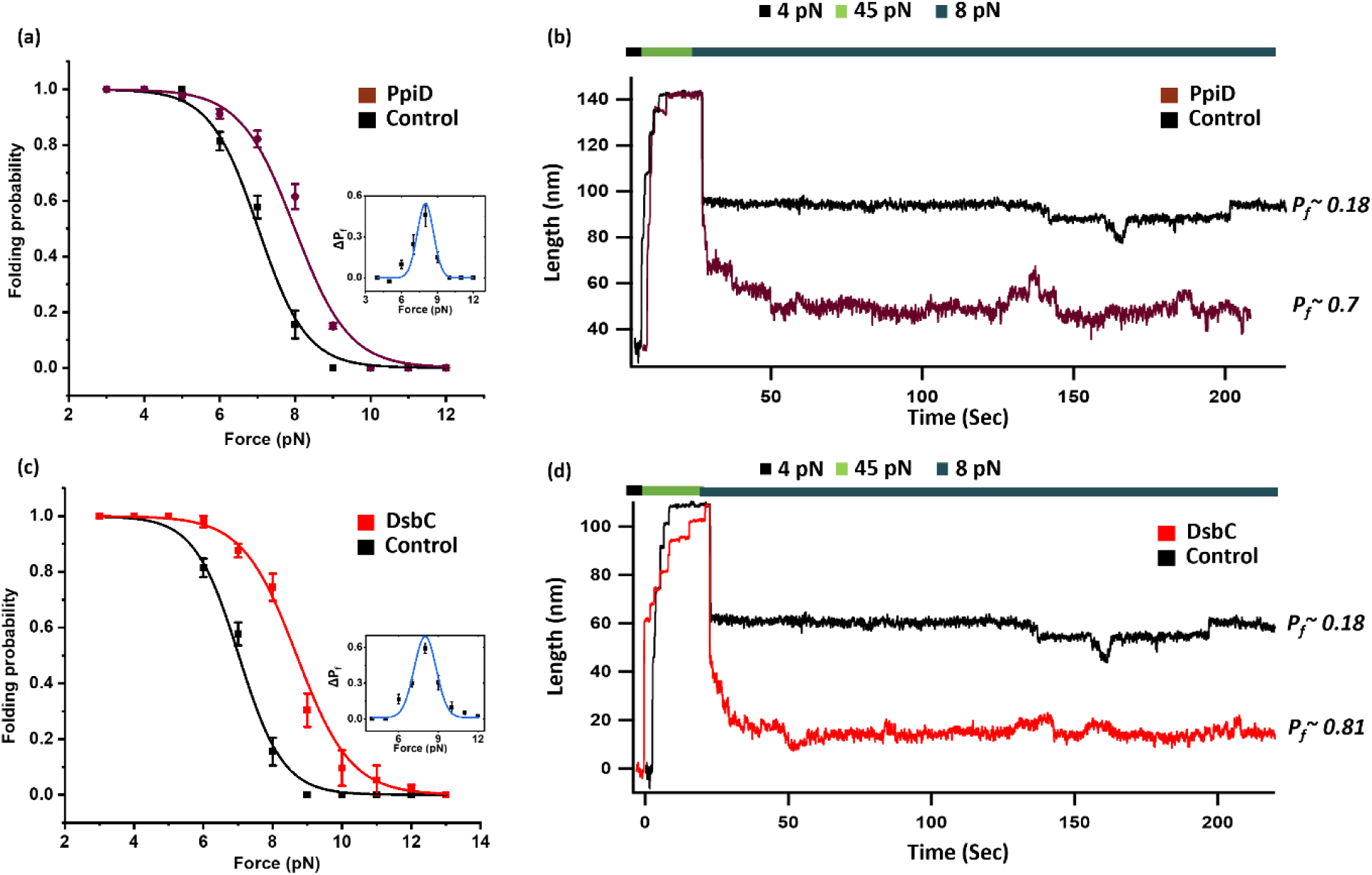
Force-clamp experiment detects the chaperone-mediated change in the folding probability of protein L: **(A)** Folding probability has been plotted as a function of the force in the presence (maroon) and absence (control, black) of PpiD. PpiD shifts the midpoint folding probability from 7 to 7.98 pN indicating the mechanical chaperone activity of PpiD towards a higher force region. Relative increment has been shown in the inset. Each data points represents >3 individual molecules and observed for >250s.. Error bars are the standard error of the mean (S.E.M.). **(B)** A representative trace for protein L in the absence and presence of PpiD demonstrates the effect of PpiD on the folding probability of the substrate protein. After unfolding the polyprotein at 45 pN, the protein was subjected to refold at 8 pN, where the folding probability of the protein L has been increased from 0.18 to 0.7. **(C)** In the presence of DsbC (red), the folding probability of protein L moves towards a higher force region, and the midpoint shifts from 7 to 8.69 pN. Each data points represents >3 individual molecules and observed for >250s. Relative change in the FP is shown in the inset, where the maximum increase in FP observed at 8 pN. Error bars are the standard error of the mean (S.E.M.). **(D)** Representative trace of protein L has been showed in the presence of DsbC. Here also the polyprotein was unfolded at 45 pN. The shifts in the folding probability have been observed from 0.18 to 0.8 at the refolding force of 8 pN.

We further investigated the effect of PpiD by correlating the kinetics of protein L at different force regimes. To implement that, we have compared the MFPT of refolding (Supp. Fig. 3A), in the presence (maroon) and absence (black) of 25 µM PpiD. At any force, the kinetics of protein refolding is greatly accelerated in the presence of PpiD. At 7 pN, the polyprotein requires 107±4.5 sec to refold completely, whereas with PpiD it takes 66.7±13 sec to refold. Such a mark change in the MFPT of refolding and FP implies that PpiD must interact with the unfolded state of the substrate protein and accelerate the folding kinetics. Similarly, MFPT of unfolding has been observed to increase marginally in the presence of PpiD (Supp. Fig. 3B).

Next, we are interested to know the mechanical effect of another periplasmic chaperone, associated with the Sec YEG tunnel, DsbC, a versatile periplasmic chaperone along with its disulfide isomerase activity^30,42^. We undertook a mechanical study of protein L in the presence of DsbC to shed light on its mechanical function as it also has evidence of direct interaction with periplasmic protein and in the translocation of the substrate. We performed the FP in the presence of DsbC under the force range of 4-12 pN and observed a rightward shift in the folding probability (Fig. 2C and 2D). The maximum shifts have been monitored at a particular force of 8 pN. The half-point force is also shifted from 7 pN to 8.7 pN in the presence of DsbC. We investigate the effect of DsbC at a concentration of 25 µM, as its mechanical effect on protein L is saturated at this level (Supp Fig. 4). Following the similar approach used in the case of PpiD, here we also measured the kinetics of protein L with DsbC. At a particular force, DsbC accelerates the refolding kinetics by decreasing the MFPT of refolding. For example, at 7 pN DsbC takes 53±19 sec to refold while it took 107±4.5 sec to refold in the absence of DsbC (Supp. Fig. 5A). Likewise, MFPT of unfolding was also shown to increase in the presence of DsbC indicating the stabilization of the folded state (Supp. Fig. 5B).

### The holdase activity of Spy and Skp under mechanical stress

Our investigations have primarily focused on SecYEG tunnel associated chaperones; however, to further explore the roles of freely moving periplasmic chaperones, we extended our study to investigate the mechanical roles of the chaperones, Spy and Skp, not associated with the tunnel. Spy, an ATP-independent periplasmic chaperone, can interact with the substrate protein in both non-native and native states, but prior to interacting with the non-native state^33^. Spy interacts with its positively charged residue, having a cradle-shaped binding site for interacting with the unfolded substrate protein^43,20^.

We observed that Spy shifts the FP of the client towards the lower force region and the midpoint force also decreases from 7 pN to 6.04 pN (Fig. 3A and 3B). From the kinetics study, it has been observed that Spy retards the refolding kinetics and accelerates the unfolding kinetics (Supp. Fig. 6). These results strongly implicated that Spy exerts robust stabilization effect on the unfolded state of protein L.

**Figure 3:**
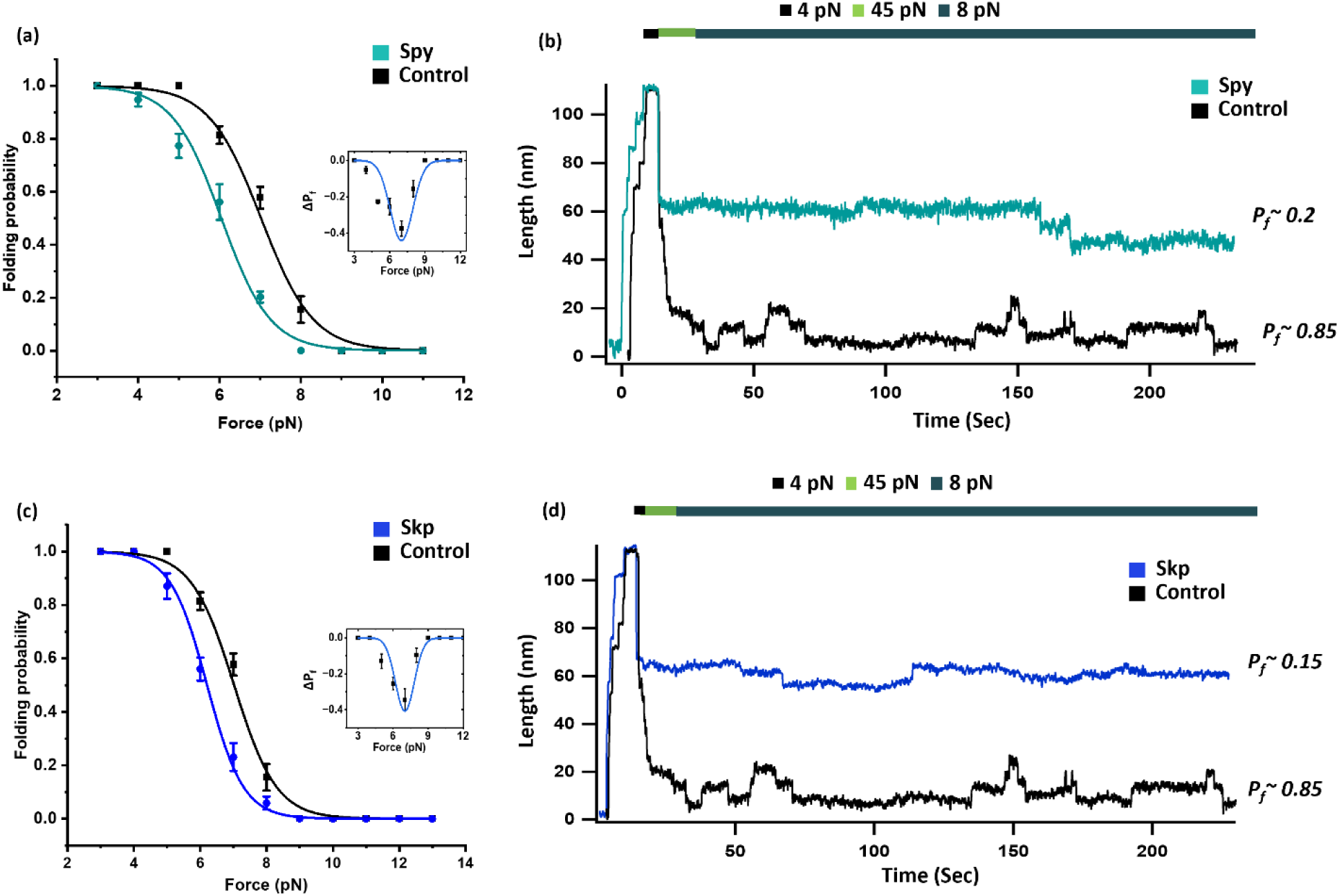
Force-clamp experiment detects the chaperone-mediated change in the folding probability of protein L: **(A)** The leftward shift of the curve in the presence of Skp (blue) has been observed which indicates that Skp reduces the folding probability towards a lower force regime. Relative change has been shown in the inset, where the maximum decrease in the FP has been observe at 7 pN. Each data points represents >3 individual molecules and observed for >250s. Error bars are the standard error of the mean (S.E.M.). **(B)** Representative trace of protein L in the presence and absence of Skp highlights its impact on the folding probability of protein L at a particular refolding force of 8 pN, where the folding probability of protein L is increased from 0.15 to 0.85. **(C)** Similar to Skp, Spy (aqua blue) also decreases the folding probability at a particular force and the midpoint force of FP shifts from 7 pN to 6 pN. Relative change is shown in the inset, with the maximum decrease in FP observed at 7 pN. Each data points represents >3 individual molecules and observed for > 250 s. Error bars are the standard error of the mean (S.E.M.). **(D** A representative trace for protein L has been recorded in the absence and presence of Spy. The folding probability of protein L decreased significantly from 0.85 to 0.2 in the presence of spy at a refolding force of 8 pN suggesting its roles as an holdase under mechanical force.

Next, we moved to another periplasmic chaperone Skp, which helps to transport the client protein in its unfolded state over the periplasm towards the outer membrane^21,44^. Here we found that similar to Spy, a leftward shift in the folding probability has been observed in the presence of Skp (Fig. 3C and 3D) which in turn retards the refolding dynamics. We further measured the MFPT of refolding which implies that at a particular force, the MFPT of refolding is marginally higher in the presence of Skp, implying slower refolding kinetics (Supp. Fig. 7) A marginal change in the unfolding kinetics has been observed. This result implies the holdase activity of Skp, characterized by its ability to stabilize the unfolded state of the substrate proteins.

Expanding our exploration, we investigated the roles of chaperone in a concentration-dependent manner and observed that beyond certain concentration, the effects of Skp and Spy on the folding dynamics become saturated (Supp. Fig. 8 and 9). The change in the folding probability with increasing the concentration of the chaperone was fitted with the hill equation, which implies that the effect of the chaperone on folding probability becomes maximum at 10 and 25 µM for Skp and Spy respectively.

### Effect on folding probability on the folding dynamics of talin

To generalize our result, we systematically investigate the effect on the folding dynamics of another two-state model substrate talin^26^. Talin is a two-state mechanosensitive protein, structurally different from protein L, in-vivo function under mechanical force^26,45,46^. We used R3 IVVI domains of talin, an alpha helix domain-containing protein, district from protein L both structurally and functionally. Here we have measured the folding probability of talin by the force clamp method and calculated the midpoint force in the presence and absence of chaperones (Fig. 4). The midpoint force for talin was observed at 7.9 pN in the absence of any chaperone. However, in the presence of DsbC and PpiD (Fig. 4A and 4B), a notable upward shift in the half-point force was showing the mechanical foldase activity of those chaperones. Further examination on the folding activity of talin in the presence of periplasmic chaperones Skp and Spy (Fig. 4C and 4D) demonstrated a downward shift in the midpoint force from 7.9 pN to 6.27 and 7 pN respectively. These results illustrate the various molecular roles of chaperones in regulating protein folding under mechanical stress and reveal the intricate interaction between chaperones and the substrate protein folding dynamics.

**Figure 4:**
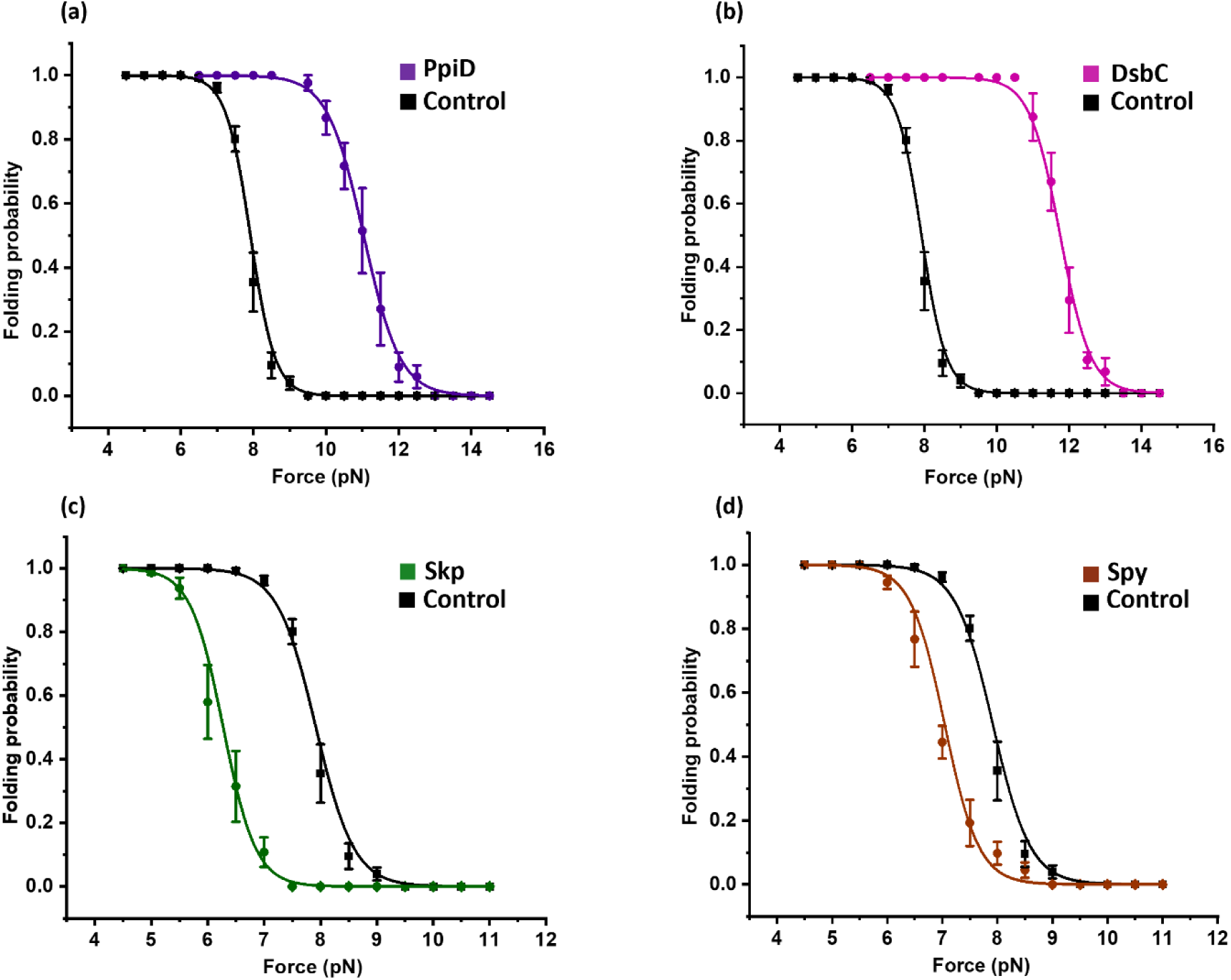
Comparison in the folding probability of talin modulated by periplasmic chaperones: **(A)** Folding probability of talin shifted in favour of the folded state in the presence of PpiD (purple) and the half point force increases from 7.91 to 11 pN. **(B)** In the presence of DsbC (pink), the FP of talin moves towards the rightward side of the curve indicating the increases in FP toward higher force region. **(C)** Skp (green) shifts the folding probability towards the lower force region and the midpoint force from 7.91 to 6.27 pN. **(D)** Spy (brown) also declines the folding probability towards lower force region by shifting the midpoint force from 7.91 to 7 pN. Each data points represents >3 individual molecules. Error bars are SEM.

### Mechanical energy provided by substrate protein assisted by periplasmic chaperone

The development of force-clamp technology demonstrated that, upon quenching, the unfolded protein gets refolded by shortening the length towards the folded state and generating mechanical work against the pulling force^14,47,48^. The total amount of mechanical work done by the system or on the system can be quantified by multiplying the step size with the applied force as described before by Eckles et al^39^. The distribution curve shows that (Fig. 5) the mechanical work output is highest at higher force region. However, at higher force, the probability of folding is lesser, so it is more convenient to get the expected value of work done by multiplying it by folding probability. We therefore calculated the mechanical work done in the presence of chaperones for two different substrate protein, protein L and talin. Therefore, due to the change in the folding probability we have observed the impact on the mechanical work done on the substrate protein in the presence of different chaperones. This distribution curve (Fig. 5A) shows that protein L can generate maximum 29.1 zJ of work at 7 pN force. Whereas in the presence of a foldase chaperone like PpiD, this peak value increases up to 45.5 zJ, the same increment has also been observed in the case of other foldase DsbC. However, for holdase chaperones like Skp or Spy, generating a lower amount of mechanical work outputs as compared to protein L. Fig 5B shows that talin can generate mechanical work of 139.3 zJ at a peak force of 7 pN, but in the presence of PpiD, it generates 166.8 zJ of work at 10 pN. In a similar way, the maximum mechanical work is reduced for the holdase chaperone. This result suggests that foldase chaperone PpiD and DsbC deliver extra mechanical work to recover the client protein from the Sec YEG tunnel whereas holdase chaperones deliver a lower amount of energy for protein translocation.

**Figure 5:**
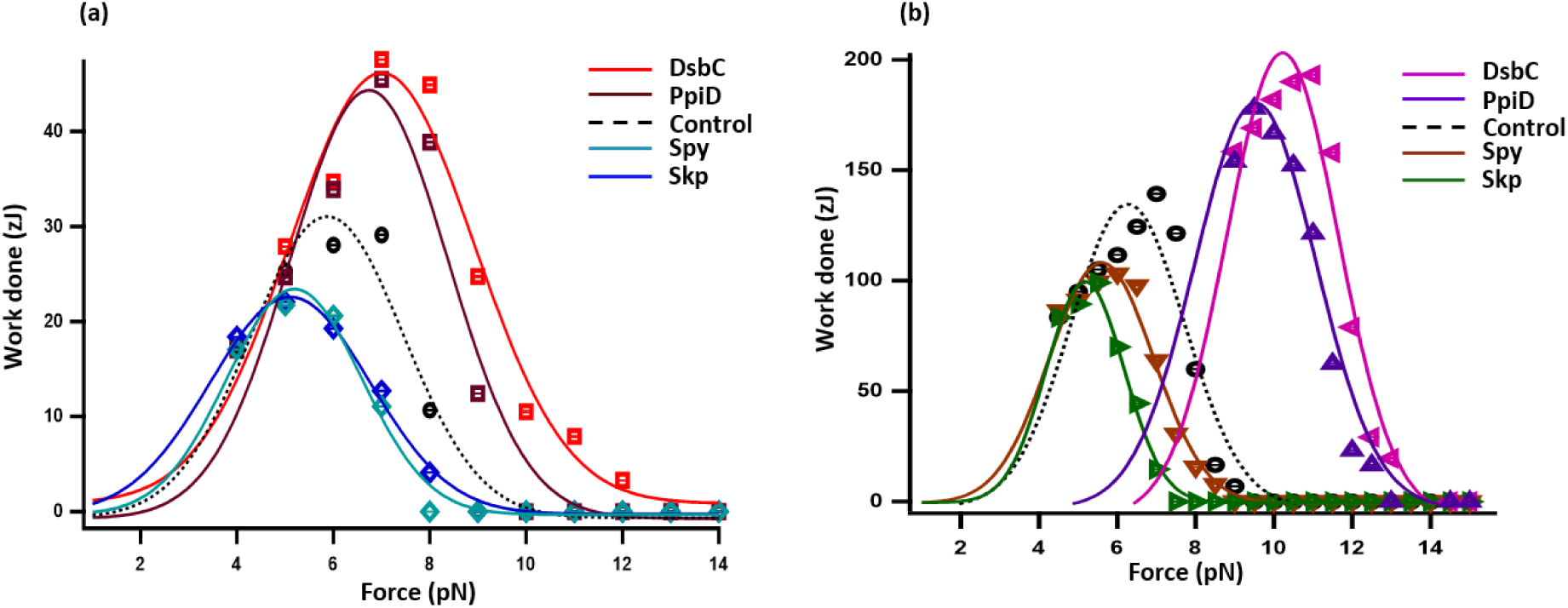
Mechanical energy delivered by different chaperones during protein folding at equilibrium: **(A)** Mechanical work done by protein L has been calculated by multiplying the folding probability with force and step size at force-induced equilibrium condition. Foldase chaperones PpiD and DsbC cause gigantic effects on the mechanical work done delivered by protein folding by increasing the mechanical energy, whereas Skp and Spy reduce the mechanical energy and generate lower mechanical work. **(B)** Mechanical work delivered by talin has been calculated in a similar fashion. PpiD and DsbC generate highest mechanical work of 166.8 and 193.16 zJ at 10 and 11 pN respectively. Holdase chaperone Skp and Spy generates lower mechanical energy by 99 and 103.2 zJ at 5.5 and 6 pN. Error bars are SEM.

## Discussion

Advanced technologies in understanding the molecular mechanics of the periplasmic chaperone involved in outer membrane protein biogenesis have been studied widely in recent years^20,22,44,49^. However, there is still a lack of direct quantitative evidence of how force can control the molecular mechanism of the folding dynamics of the substrate proteins at the periplasm. Here we provided a single molecule force spectroscopic technique to directly explore how chaperones-client interaction could be modulated by mechanical load. Our data suggest a potential mechanism for periplasmic chaperones, shedding light on how they may decrease ATP consumption by performing additional mechanical work within the tunnel.

Our present study provides an in-depth mechanistic detail of how tunnel-associated periplasmic chaperones initiate the refolding of the client protein in the presence of force. Recently, it was shown that trigger factor (TF), associated with ribosomal tunnel engages with the newly synthesized polyprotein and acts as a mechanical foldase by assisting the protein to refold as they emerge from the ribosome^16^. Ulrich et. al found some structural homologous domains of cytosolic chaperone trigger factor with PpiD, a periplasmic chaperone associated with the SecYEG tunnel^50,51^. Previously, it was also shown that PpiD helps to facilitate the newly synthesized protein from the SecYEG channel, similar to Skp and shown to interact with the model peptides to prevent aggregation of denatured protein^29,32,41^. Here we have found out that PpiD interacts with the client under force and acts as mechanical foldase by shifting the FP to the higher force region. Next, we sought to investigate the chaperone activity of another tunnel-associated periplasmic chaperone DsbC having the hydrophobic cleft that has been postulated to be responsible for the hydrophobic interaction with the client protein^52,53^. Our finding reveals that DsbC interacts with the substrate under force, potentially enhancing the folding probability as well as refolding kinetics, implicating its foldase activity through the binding energy. According to the Brownian ratchet mechanism during the translocation of the substrate in the Sec machinery, the backsliding of the substrate towards the cytoplasm may be prevented by the binding of the periplasmic chaperone, as discussed previously by Allen et al^54^. Thus, our result presents a in depth mechanism of the periplasmic chaperones during the translocation of their substrates through the SecYEG pathway to the periplasmic side of the inner membrane, which reconciles the phenomena of the ratchet mechanism.

Furthermore, we wonder to investigate how the involvement of force in the chaperone-substrate interaction could be affected the ATPase activity of the SecA. Previously it was reported that SecA translocate the unfolded polypeptide through the SecYEG machinery by using the energy of ATP hydrolysis where chemical energy of ATP can perform mechanical work^55^. Notably, SecA can translocate 20 amino acids through the translocation pore, with the hydrolysis of one ATP molecule^56^. Thus, to translocate one protein L B1 chain having 62 amino acids, SecA requires to hydrolysis 3 molecules of ATP, which outputs 150 zJ (3 ATP × 100 zJ per ATP × 50% efficiency) of mechanical work^55–58^. Recent single molecule force spectroscopic study reveals that SecA motor generates a mechanical force of ∼10 pN during translocation^10^. Here, we also have observed the substrate refolding within 4-12 pN range for calculating the mechanical work delivered by the periplasmic chaperones. In our comprehensive analysis, we obtained that the foldase chaperone DsbC and PpiD significantly enhances the efficiency of the mechanical work capacity of substrate protein. As depicted in Fig. 5A, protein L generates a maximum work of 29.1 zJ, which increases to 45.5 zJ (a 1.6-fold enhancement) in the presence of PpiD. The extra mechanical work delivered by the foldase chaperone, could help to translocate the polyprotein substrate by lowering the total ATP consumption. A similar result also observed in case of talin R3 (124 amino acids) which increases the work done by 166 zJ in the presence of PpiD. A comparable effect was observed with DsbC, underscoring the critical role of mechanical foldase chaperones in promoting protein refolding under force. In contrast to tunnel-associated periplasmic chaperones (PpiD or DsbC), freely moving periplasmic chaperones (Skp and Spy) exhibit holdase activity, stabilizing the unfolded state (Fig. 6). This is likely because, unlike their tunnel-associated counterparts, they function without force, holding the protein in an unfolded state to prevent misfolding during transport.

**Figure 6:**
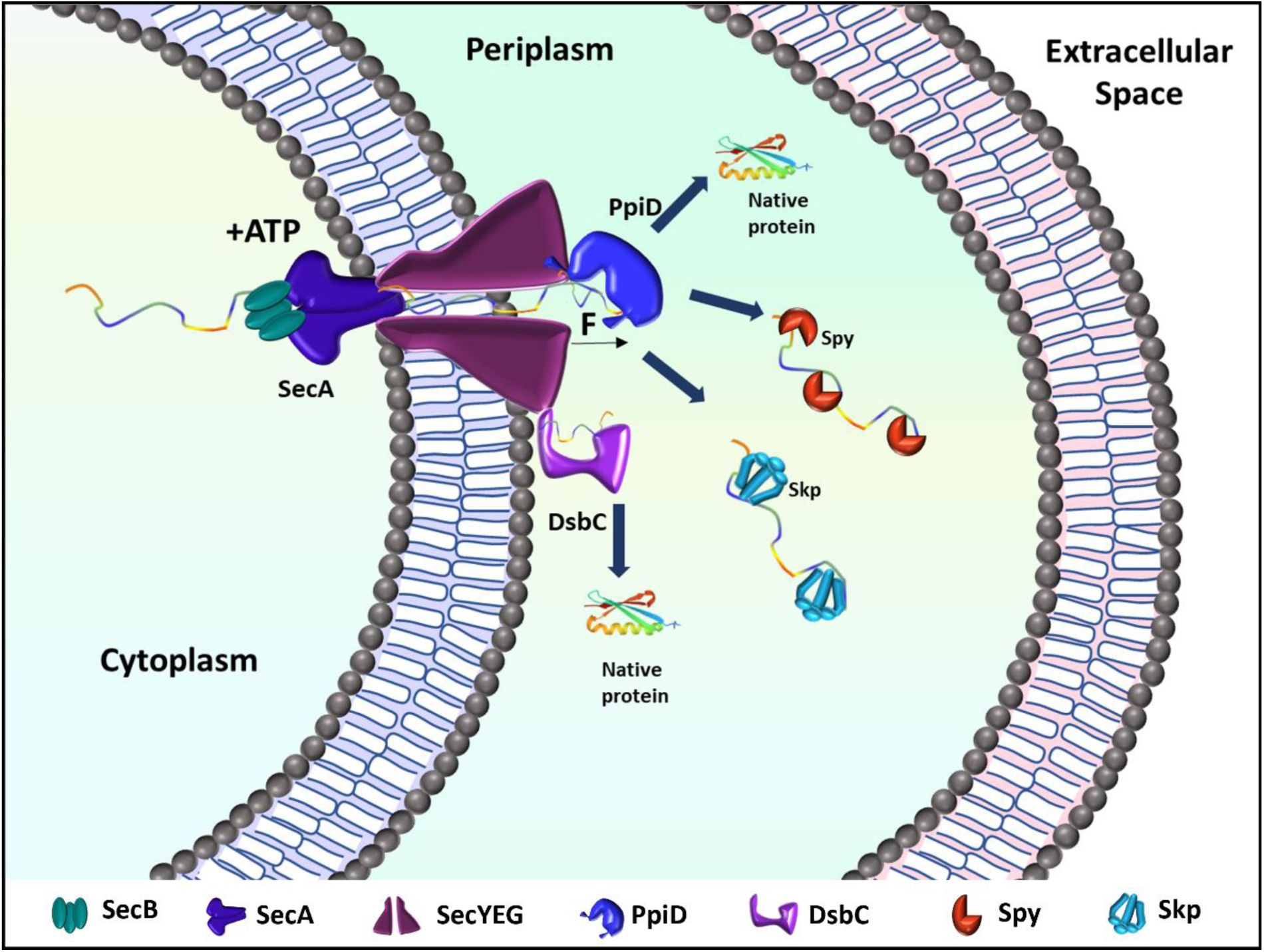
Illustration of force-regulated folding pathway in the presence of periplasm chaperone: The unfolded polypeptide can be translocated through Sec YEG translocon pore where PpiD and DsbC are the prior chaperones which interacts with the nascent polypeptide and increases the refolding yield or makes translocation faster for the unfolded polypeptide chain under mechanical stress. In another pathway the unfolded nascent chain can interact with Skp and Spy, act as mechanical holdase that helps to translocate the unfolded polypeptide through the periplasm to the extracellular space and prevent them from misfolding.

## Analysis

All the data analysis and acquisition and analysis were performed with Origin pro and Igor Pro 8.0 software (Wavemeters) software.

## Author contribution

S.H. and D.C. designed the project. D.C. and M.B. performed the experiment. D.C. analysed the data and wrote the manuscript.

## Acknowledgements

We thank the department of Chemistry, Ashoka University, S.N.Bose National Centre for Basic Science and Technical Research Center of S.N.Bose National Centre for Basic Science for the support and funding.

